# ‘Superbugs: A Pop-up Science Shop’: Increasing public awareness and knowledge of antimicrobial resistance by taking science to the city

**DOI:** 10.1101/2021.06.16.443502

**Authors:** Jonathan M. Tyrrell, Christie S. Conlon, Ali F. Aboklaish, Sarah Hatch, Carl Smith, Jordan Mathias, Kathryn Thomson, Matthias Eberl

**Author notes:** Corresponding author: Dr Jonathan M. Tyrrell [ ].

## Abstract

‘Superbugs: A Pop-up Science Shop’ was a public engagement event in the school summer holidays of 2019, organised by members of Cardiff University’s School of Medicine. We transformed an empty retail unit in the centre of Wales’ largest shopping centre into an interactive and immersive microbiology experience. We aimed to facilitate opportunities for two-way dialogue to impart positive impact on the awareness of antibiotic resistance, whilst concurrently evaluating the efficacy of an engagement strategy focused on the utilisation of public spaces to attract public demographics diverse to those who would normally engage with STEM outreach.

Over the course of 14 days, we welcomed 6,566 visitors, with 67% attending as part of the natural footfall of the shopping centre. We created 1,625 young Antibiotic Resistance Champions, located in over 200 schools. Through a multi-lateral evaluation strategy, we were able to collect quantitative and qualitative feedback on the success of our delivery model, and the impact on our stakeholders. Herein, we will discuss the evolution of ‘Superbugs’ from concept, planning and design, to the logistics of delivering an engagement event of this scale. We will focus in particular on the learning outcomes of the project, and how this will shape the future of our ‘Superbugs’ project, and engagement events beyond.

**Key Messages:**

1. Creating a multi-disciplinary core team is essential to the success of large-scale engagement events as well as the support and development of large numbers of colleagues/volunteers
2. Utilising themes of exhibition and gameplay alongside strong fear-empowerment messages is an impactful way to confer positive influence and behaviour around antimicrobial resistance (AMR) and the use of antibiotics
3. ‘Pop-up shop’ is an effective mode of delivery to capture diverse public demographics far beyond those who would traditionally engage with scientific outreach and science engagement.

## 1. Introduction

Antimicrobial resistance (AMR) is among the most significant threats to global public health, food security, and development (https://www.who.int/news-room/fact-sheets/detail/antibiotic-resistance). Infections resistant to multiple classes of antibiotics compromise patient treatment and carry significant economic burden. The spread of such infections is exacerbated by socioeconomic factors including, but not limited to, the inappropriate use of antibiotics at a local, national and international level. Given the everyday implication to public health, social education and participation is paramount to the success of strategies to control the concerning rise of AMR worldwide. Society has a pivotal role to play through improved antibiotic stewardship, improvement and maintenance of good hygiene, vaccination and the election and accountability of policy makers. Management of major AMR pathogens such as methicillin-resistant *Staphylococcus aureus* (MRSA) and *Clostridium difficile* has shown that public-focused campaigns around infection control and antibiotic stewardship can indeed result in reduction of infection rates (Ashiru-Oredope *et al*., 2012; Duerden *et al*., 2015). However, without adequate insight and basic understanding of infection control and antibiotic resistance on behalf of society, we cannot hope for success in programmes and initiatives aimed at combatting the current crisis. The WHO Global Action Plan on Antimicrobial Resistance outlined Objective 1 of their strategic plan as ‘Improve awareness and understanding of antimicrobial resistance through effective education and training’ (WHO, 2015a), testimony to the importance of public participation and understanding. On 24^th^ January 2019, the UK government identified ‘Engage the public on AMR’ as one of their nine ambitions as part of their 20-year vision for antimicrobial resistance (https://www.gov.uk/government/publications/uk-20-year-vision-for-antimicrobial-resistance).

In conjunction with their action plan, the WHO published a 12-country snapshot survey, the ‘Antibiotic Resistance: Multi-country Public Awareness Survey’ (WHO, 2015b). Of those surveyed, 76% incorrectly identified the statement ‘Antibiotic resistance occurs when your body becomes resistant to antibiotics and they no longer work as well’ as true. 64% incorrectly thought that a common cold and the flu can be treated with antibiotics. Perhaps most concerningly, only 18% of participants disagreed with the statement ‘There is not much people like me can do to stop antibiotic resistance’. There are economic, geographical and educational caveats to many of the results presented in the report, but the picture painted is a stark one. Furthermore, this is not an issue limited to the lay public. Dyar *et al*., (2018) reported that, whilst 100% of healthcare students (predominately medicine, pharmacy, dentistry and veterinary medicine) correctly stated that bacteria could become resistant to antibiotics, more than 40% also incorrectly believed that humans and animals could also.

It is with active members of the AMR and infection research community, such as ourselves in the ‘Superbugs’ team, where responsibility must lie; there is clearly inadequate and inefficacious communication of the AMR message, the research being carried out, and the role society has to play, at a time where it has never been more pertinent. Not only due to the grave consequences the global AMR crisis does, and will, continue to have on mortality and morbidity, and economic implications, but also to avoid the assuaging of impact of research within the field. It is through these frustrations, and a desire to contribute to the readdressing of the balance, that our project, ‘Superbugs: A Pop-Up Science Shop!’ was born.

## 2. Aims of Manuscript

‘Superbugs: A Pop-Up Science Shop!’ was a proof-of-concept event, held in the school summer holidays between 29^th^ July and 11^th^ August 2019, open for all 14 days in a row, 9am-5pm. The event aimed to combine the philosophies of public engagement with research, school outreach and science exhibition to deliver an interactive and immersive experience in a public space, outside of traditional scientific and academic environments. The event utilised a vacated retail unit (approximately 2400 ft^2^ in size) in the public space of St David’s (StD), the largest shopping centre in Wales and one of the busiest in the whole of the UK and transformed the retail unit into a professionally designed interactive research experience involving laboratory activities, games, artworks, information resources and competitions surrounding the topics of bacteria, antibiotics and drug-resistant infections.

Herein, we will explore in detail the process of conception, planning and delivery of the ‘Superbugs’ event. Further to this, we will present much of the quantifiable output generated by our stakeholders and the project team, and how this learning may be of positive impact for (AMR) researchers, public engagement professionals and all organisations with a public engagement agenda

## 3. Planning of Project

### 3.1 Conception

The concept of ‘Superbugs: A pop-up Science Shop!’ originated from two earlier key events. The first was the organisation of ‘Superbugs: The End of Modern Medicine’, a public engagement evening held at Techniquest, an educational science and technology museum in Cardiff (https://www.cardiff.ac.uk/_data/assets/pdf_file/0007/1270852/Cardiff-University-ReMEDy-Newsletter-Edition-29_e_web.pdf). Through feedback collected from visitors, it was here that the potential to deliver meaningful AMR messages through the medium of exhibition, presentations, games and interactive laboratory tasks was first explored, which laid the foundation for the content developed for the later pop-up science shop. The second event was participation in the Cardiff University BLS (Biomedical & Life Sciences) Public Engagement Development Programme, through which we were introduced to the ‘Pop-Up Science’ report (Figure 1), and the potential of such a delivery model. Research by the British Science Association and King’s College London has suggested that up to 76% of UK adults (defined as 16+ years olds), accounting for approximately 49 million people in total, do not participate in scientific outreach, either due to lack of interest or effort to seek out such opportunities. This highlights the space to further evaluate the impact of scientific engagement carried out in public areas, and engagement of the ‘general public’.

**Figure 1:**
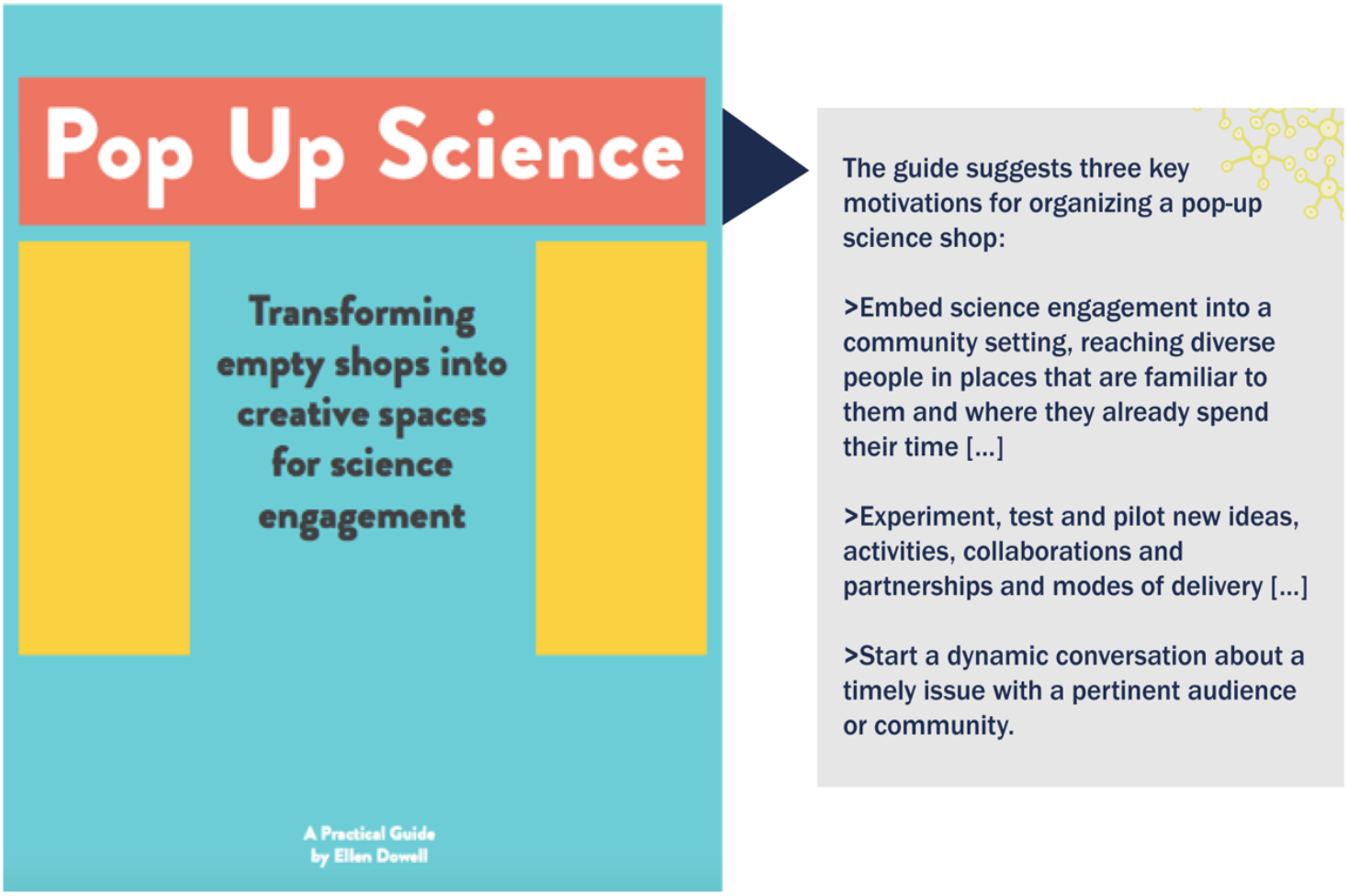
Key applicable points taken from ‘Pop Up Science-A Practical guide’ published in 2017 by Ellen Dowell from the National Heart & Lung Institute at Imperial College London Heart & Lung Institute at Imperial College London. (https://www.imperial.ac.uk/nhli/interact/public-engagement/our-projects/pop-up-science/)

Carr *et al*., (1992), in their text ‘Public Spaces’ outlined five important requirements that influenced the appeal of public spaces on communities: (i) comfort, (ii) relaxation, (iii) passive engagement, (iv) active engagement, and (v) discovery. It was certainly the latter two, the need for mental and physical challenge (active engagement) and offering the chance to evolve new ideas and interests in unfamiliar topics (discovery), that were at the core of what we hoped to achieve in this project. As Carr *et al*., so succinctly put it, ‘enable the users’ interest to endure’.

### 3.2 Application for Funding

Our plans were collated into an application to the Wellcome Trust Institutional Strategic Support Fund (ISSF3) on the theme of Public Engagement Proof-of Concept funding, with the proposed AMR pop-up science shop as a novel delivery model that had not been utilised before at Cardiff University, nor to our knowledge more widely in the area of AMR. The overall aim for the proposed project was to increase the awareness and knowledge of the general public of the microbial world, infection biology and the increasing threat of antibiotic resistance to global public health. Additionally, we hypothesised that by taking the event into public spaces, we could evaluate the efficacy of attracting the attention of all public demographics, and not only those that would traditionally attend scientific engagement events (Figure 2).

**Figure 2:**
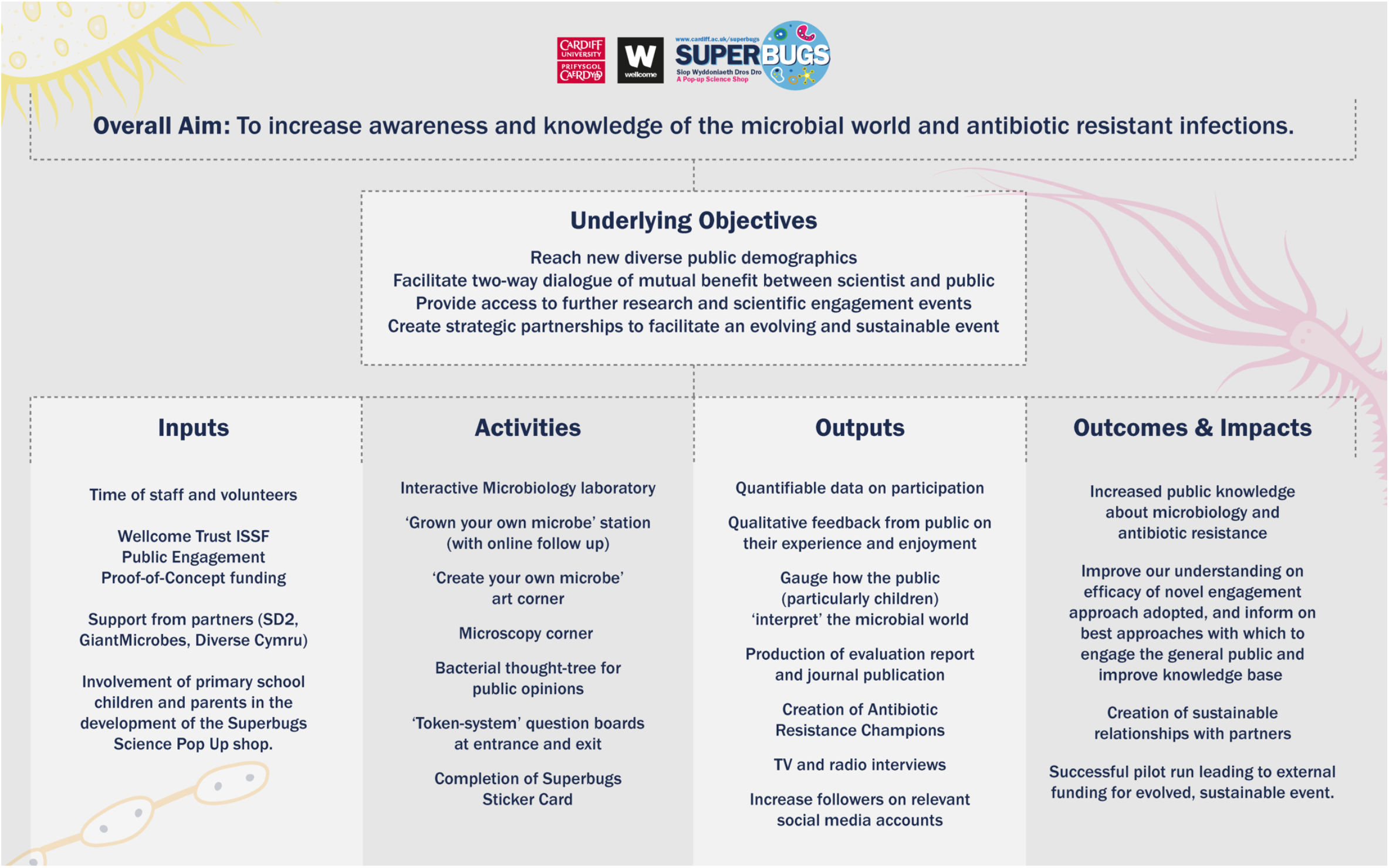
Logic Model of ‘Superbugs: A Pop-Up Science Shop’.

In order to achieve this, we defined a number of underlying objectives;

i. To reach new and diverse public demographics by taking science to the people
ii. To facilitate two-way dialogue of mutual benefit between scientists and the public
iii. To provide the public access to information about local and international AMR-related research, how it impacts their lives, and further scientific-public engagement opportunities
iv. To create strategic relationships between Cardiff University, the public, and supporting partners.

In September 2018, we were successfully awarded Public Engagement Proof of Concept funding and secured contingency funding and support from the Systems Immunity Research Institute at Cardiff University, a department carrying out world class research in AMR, infection and immunity, where most of the funded team’s scientists were based at the time. Cardiff University is the leading biomedical research institution in Wales as evidenced by ranking 5^th^ overall in the UK and 8^th^ in Clinical Medicine in the Research Excellence Framework (REF) 2014 – the most recent nationwide impact evaluation assessing the research at all UK higher education institutions– with its world leading basic and clinical research on AMR, infection and immunity as a core strength.

## 4. Delivering the Project

### 4.1 Multi-disciplinary team and Strategic Partners

From an early stage it was identified that a multi-skilled team from various backgrounds was vital to delivering the project successfully. A core team was put together from across the School of Medicine including AMR research scientists, public engagement champions and professionals, and a graphic designer. The benefit of this to the project was not only the wide spectrum of talents and knowledge, but also the network of contacts brought to the table by each core team member. In addition to this, two early career researchers (ECRs) were recruited to work with us in designing the AMR-related content, of which the shop would comprise.

Equally as important to the success of the project were the strategic partners with whom we were able to engage.

- <u>St David’s (StD)</u>: The largest shopping centre in Wales based in the heart of the Welsh Capital, Cardiff, with a footfall of approximately 750,000 per week. Management staff from StD were involved in very early discussions on the viability of providing an empty unit within the centre and provided support to our funding application along those grounds. unit within the centre and provided support to our funding application along those grounds.
- <u>Morgans Consult</u>: Local signs and brand implementation specialists (https://morgansconsult.com/). Funding freed up by StD was invested into further professional implantation of both the interior and exterior of the pop-up shop. Morgans Consult worked closely with the team to design and fit an exhibition bespoke for the retail space we inherited from StD.
- <u>Diverse Cymru</u>: The only charity in Wales that focuses on all protected characteristics, to challenge discrimination and reduce inequality. Diverse Cymru provided us with an independent review, evaluating the performance of ‘Superbugs’ in meeting the needs of and appealing to stakeholders with protected characteristics.
- <u>British Society for Immunology</u>: Provided enthusiastic support for the original application and project, alongside the promise of promotional and educational material for distribution at the event.

### 4.2 Focus Group for Target Stakeholders

To inform the planning stages of the event, a focus group was organised for key target stakeholders: families with children of Key Stage 2 (7-11 years old) and Key Stage 3 (11-14 years old). The event was held after school hours at Rhiwbina Primary School in Cardiff and was attended by seven families recruited through an invitation forwarded onto parents by the Headteacher of the school. Children were split up into two groups and taken through a number of sample activities akin to what we hoped to deliver at the future event. Concurrently, we facilitated a discussion with the parent on various aspects of the project, before eventually coming together to gather further feedback from the children. Table 1 below summarises the key outputs taken form this focus group.

**Table 1:**
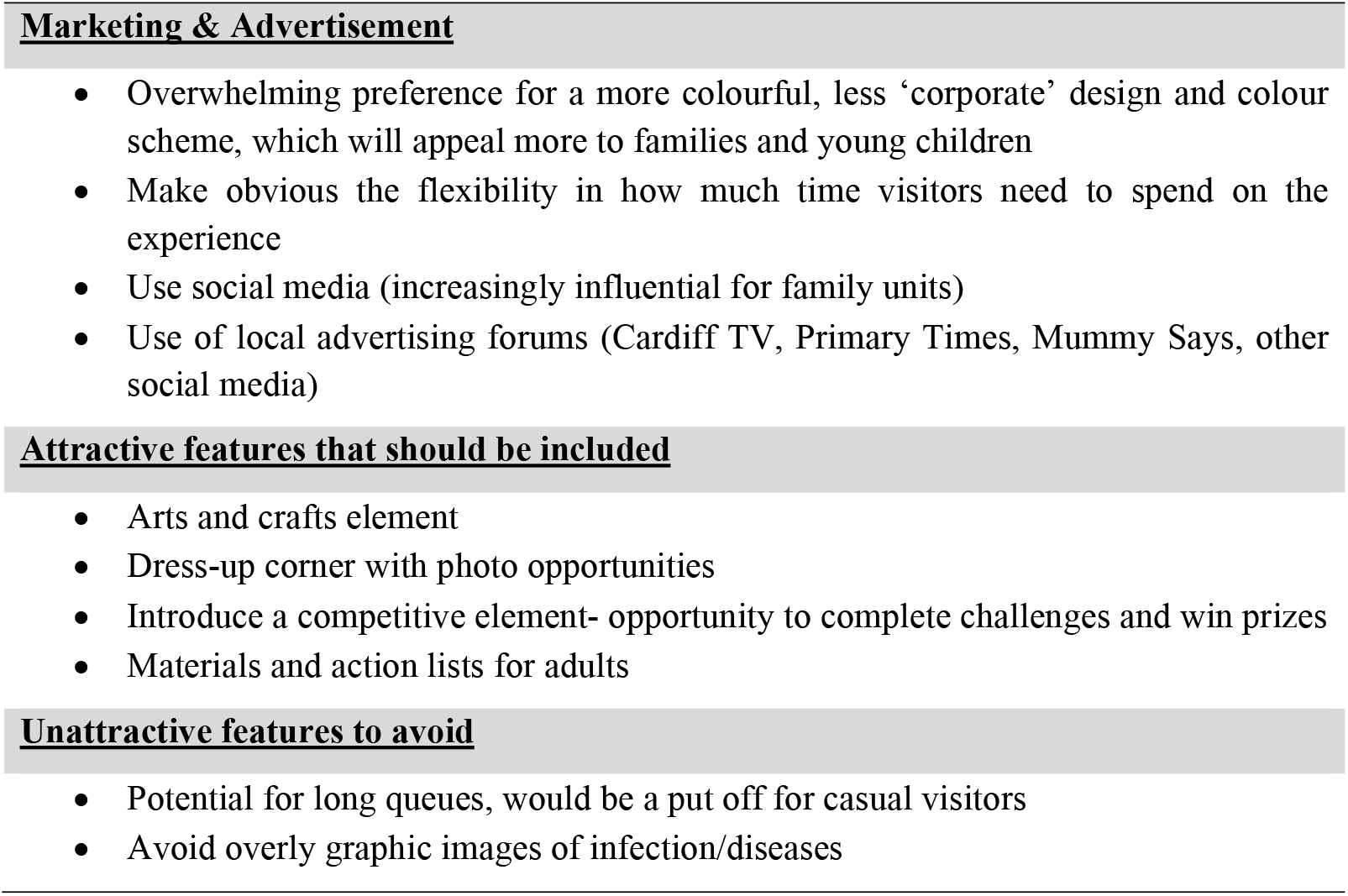
Significant output from the Stakeholder Focus Group. Discussion points taken from our Focus Group session that played a key role in the evolution and direction of the project.

This approach proved to be an invaluable step in the evolution of ‘Superbugs’, not only by informing the final design schemes of the shop, but by also confirming that our planned activities were engaging not only to our primary target stakeholders, but also by those younger and older. To quote one inquisitive four-year old after attending the focus group, ‘looking at the little things in the microscope was my favourite’.

## 5. The Content of ‘Superbugs’

### 5.1 General Design Concept & the Importance of Exterior Design

A fundamental aim of ‘Superbugs’ was to evaluate the effectiveness and success of a pop-up shop in engaging lay stakeholders of all demographics, not limited to those who would traditionally seek out STEM-related events. Attracting the natural footfall in the public location of the shop was thus imperative, and the investment of the project’s time and finance into the exterior design of the shop reflected this. Figure 3 shows the original vacated retail unit we inherited (previously a ‘Clarks’ shoe shop, with shelving and lighting still in place), and how the resulting ‘Superbugs’ pop-up shop looked once opened.

**Figure 3:**
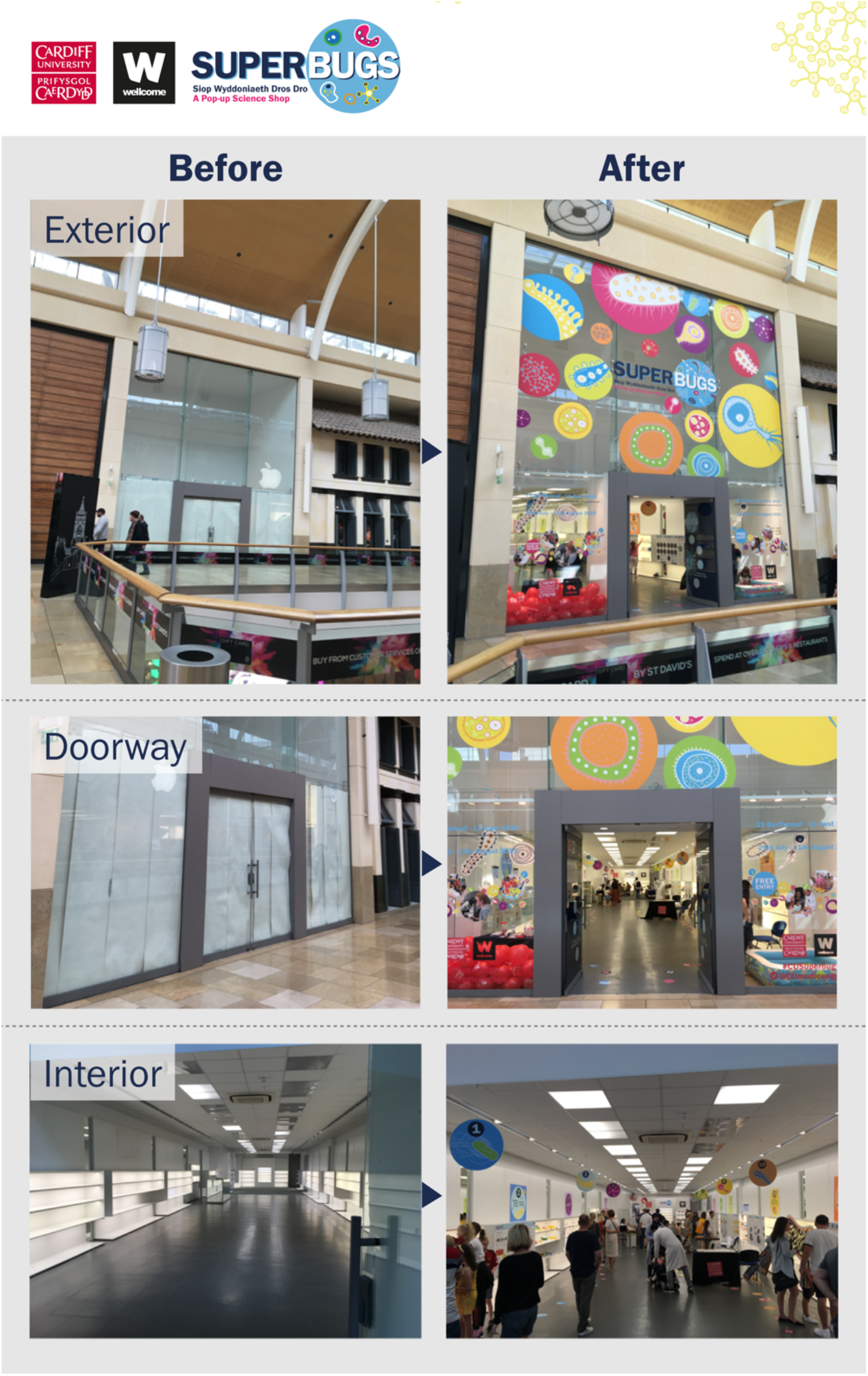
Transformation from empty retail unit to ‘Superbugs: A Pop-Up Science Shop!’. Representative photos taken before the transformation of the vacated retail in the St David’s shopping centre, and during the ‘Superbugs’ event, to illustrate the changes to the interior and exterior design of the shop.

The origin of the ‘Superbugs’ design lies in promotional material developed for the Systems Immunity Research Institute including a bus poster campaign and social media (https://www.cardiff.ac.uk/systems-immunity/about-us). Illustrations were created to represent the diversity of micro-organisms that inhabit our bodies, adding intrigue to attract further attention from the public. Acting on feedback from our focus group (Table 1), the design was overhauled for the purpose of ‘Superbugs’ using brighter colours than the original with the aim of attracting a younger audience and enlarged to create a range of super-sized ‘Superbugs’.

Strategically, the ‘Superbugs’ unit was well placed within StD. It was located directly opposite the Apple Store, within sight of other major retail units such as John Lewis. Additionally, we were situated around the corner from the major family friendly food-quarter of the mall, and close to the access point of the centre’s family crazy golf site, during the Summer holidays. All of this guaranteed a good level of natural footfall past the shop, ensuring that our large and colourful exterior design (Figure 3) had great visibility throughout the central concourse of StD.

### 5.2 Promotion of ‘Superbugs’

It is perhaps important to note that prior to the event, our promotional activities were minimal by design, to maximise the sensitivity in measuring engagement from passers-by of our public location. Firstly, we ran a minimal social media campaign, promoting our e-poster. In the week leading up to the event we ran a short promotional campaign in Primary Times Cardiff & Vale (https://www.primarytimes.co.uk/cardiff/) which involved advertising the event on social media and featuring ‘Superbugs’ in their ‘What’s On’ pages. An email with an e-poster advertising the event was sent around to secondary and primary Schools in Wales that were already in the engagement network of the School of Medicine through our ‘Science in Health’ events (https://www.cardiff.ac.uk/medicine/about-us/engagement/science-in-health). In the first few days of the live event, we carried out a leaflet drop at particular tourist and visitor hotspots around the city centre, including the nearby library, museum, and information desks.

### 5.3 Scientific Content

The Wellcome Trust ‘Reframing Resistance Report’ (https://wellcome.org/sites/default/files/reframing-resistance-framing-toolkit.pdf) identified the importance of how we approach AMR engagement; “How an issue is ‘framed’-explained and presented through specific themes and angles-can influence how it is received by an audience”. Informed by a multi-phase international research project, the report outlined a ‘Framing Toolkit’ identifying five principles in how AMR-communication and outreach should be framed; (i) frame antimicrobial resistance as undermining modern medicine, (ii) explain the fundamentals succinctly, (iii) emphasise that this is a universal issue; it affects everyone including you. (iv) focus on the here and now, (v) encourage immediate action. In co-ordinating these frames, you are likely to create communication that informs, motivates and persuades.

Roopes *et al*., (2017) previously demonstrated that a message of warning on the dangers of AMR was not successful in imparting a positive influence on either the public’s attitude or practice in requesting unnecessary antibiotics, particularly in cohorts with low AMR-awareness. Subsequent studies revealed that combining a strong fear warning with messages of empowerment for the stakeholders did in fact induce a positive response in a way that ‘fear-only’ and ‘mild-fear-plus-empowerment’ messages did not (Roope *et al*., 2020), resulting in patients being less likely to request antibiotics. The paper concluded that ‘fear could be effective in public campaigns to reduce inappropriate antibiotic use but should be combined with messages empowering patients to self-manage symptoms effectively without antibiotics’. These findings certainly reflect similar such conclusions from public campaigns in other areas of healthcare science.

We adopted such approaches when designing the content of the ‘Superbugs’ shop. We did not shy away from showing the symptoms and wider burdens of particular examples of infections, and the broader implications of AMR in the failure to treat these, which highlighted the current impact of drug-resistant infections, rather than projections or apocalyptic frames. However, alongside this was a message of positivity, detailing the many strategies by which we are fighting the AMR issue, illustrated using multiple examples relevant to the audience we were engaging. This included improvements to sanitation, agricultural use, surveillance, and the development of new drugs and vaccines, demonstrating how drug-resistant infections are a cross-cutting threat across all of society (beyond specific disease areas). Further to this we provided many sources of information on how the stakeholders can play an important role in the fight, framing the issue as solvable and providing specific calls to action e.g. complete a full course of antibiotics when prescribed. In order to be as inclusive as possible, and in line both with the recommendations of the Welsh Government and the specifications of the original grant remit, all activities and information sheets were available bilingually, in both English and Welsh.

It was decided early on in the project to avoid making any assumptions as to prior knowledge of the topic of AMR on the stakeholders, as to not compromise our overall message. As such, it was a concerted decision to begin the narrative of the event on the very basics of microbiology and bacteria that live in, on and around us, before introducing the ideas of infection and treatment. At the forefront of our messaging was an emphasis that AMR is a universal issue, and that anyone could be affected. The project included explanations of the part that human activity is playing in accelerating the issue. In this way we hoped to engage the public in AMR from an informed position on their part. Two ECRs working in the area of AMR were recruited to work alongside the core team’s scientists to form a ‘scientific content’ team, to develop the narrative and details of the exhibition. A description of the stations open to the stakeholders can be seen in Figure 4. Title cards on the wall and hanging from the ceiling, alongside bacterial ‘footprints’ indicated the suggested route around the shop, but visitors were welcomed to engage on their own terms.

**Figure 4:**
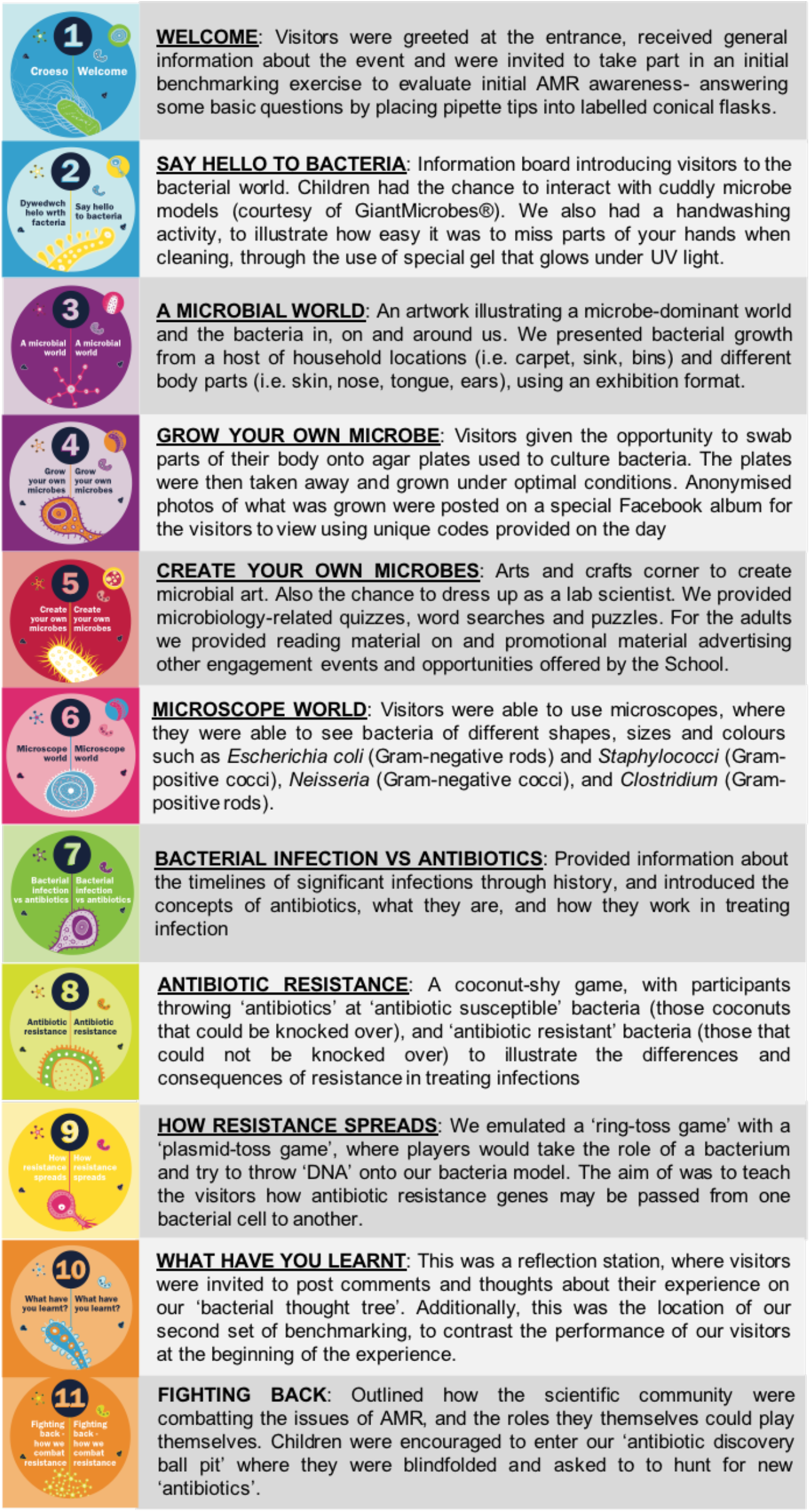
A outline of the Stations of ‘Superbugs: A Pop-up Science Shop!’

## 6. Project Output

### 6.1 Evaluation Strategy

It is widely acknowledged that rigorous and impactful evaluation of public engagement events is challenging, particularly in multi-faceted projects such as ‘Superbugs’. We not only wished to evaluate our ability to improve the awareness of AMR, but also evaluate the forum of a pop-up science shop itself as an effective public engagement delivery strategy. To achieve this, we took a multidimensional approach, collecting data in a number of varying ways to provide as detailed a dataset as possible, with reference to a common evaluation framework proposed by Reed *et al*., (2018). This framework provided a philosophical backdrop for our evaluation strategy and we will adopt a number of the principles outlined within, in future iterations of our ‘Superbugs’ events. As part of our evaluation strategy, the team reviewed the possible outcomes and impact of the novel project approach through development of a logic model (Figure 2), exploring the inputs, outputs, activities, outcomes and impacts of the project. This was adapted on numerous occasions to ensure feasibility and reach.

### 6.2 Qualitative Output

Over the course of two weeks, we welcomed a total of 6,566 visitors to ‘Superbugs’. This vastly exceeded our expectations of 1,000-1,500 per week (modelled on previous pop-up shop examples, and as such we had to adapt quickly to meet the overwhelming need for consumable elements of the shop (arts & crafts, culture plates for Station 4, questionnaires and sticker cards). Typically, visitors explored at the event for anything between 10 minutes to 1 hour.

Questionnaires were completed by 10% (n=656) of visiting parties, which is particularly impressive when taking into account that a significant number of our stakeholders were family groups with, in many cases, one parent/child completing a questionnaire on behalf of the whole family. Returning a completed questionnaire to the organisers was incentivised by entry into a twice-daily prize draw, where the winners won a bag of ‘Superbugs’-branded prizes. Station 5 (‘Create your own microbes’) produced over 500 pieces of artwork, many of which adorned the walls of the shop throughout the event and featured prominently in our social media campaign. Over 300 items of activities and reading material were taken away from Station 5 by visitors. At Station 4 (‘Grow your own microbes’) we generated 2,169 swab plates, with the subsequent social media posts garnering over 2,500 views.

Owing much to these swab photos, our social media presence gained significant interest (Figure 6). The Systems Immunity Research Institute Facebook page, where we posted the swab photos (https://www.facebook.com/SystemsImmunity), saw a 233% increase in traffic over the course of the event, with 6,000-9,000 new weekly organic impressions and 2,000 direct hits on the official ‘Superbugs’ website at Cardiff University. On Twitter, we saw over 140,000 impressions across the three main Twitter accounts promoting the event; @JTyrrell_Micro, @CUSystemsImmu and @CUMedicEngage. As the official Twitter page for the School of Medicine, @CUMedicEngage saw a 20% increase in followers over the course of the ‘Superbugs’ event. The Twitter hashtag #CUSuperbugs evidences the positive experiences of individuals accessing the AMR public engagement with research interactive science shop and provides a visual and storytelling narrative of the public experience.

**Figure 5:**
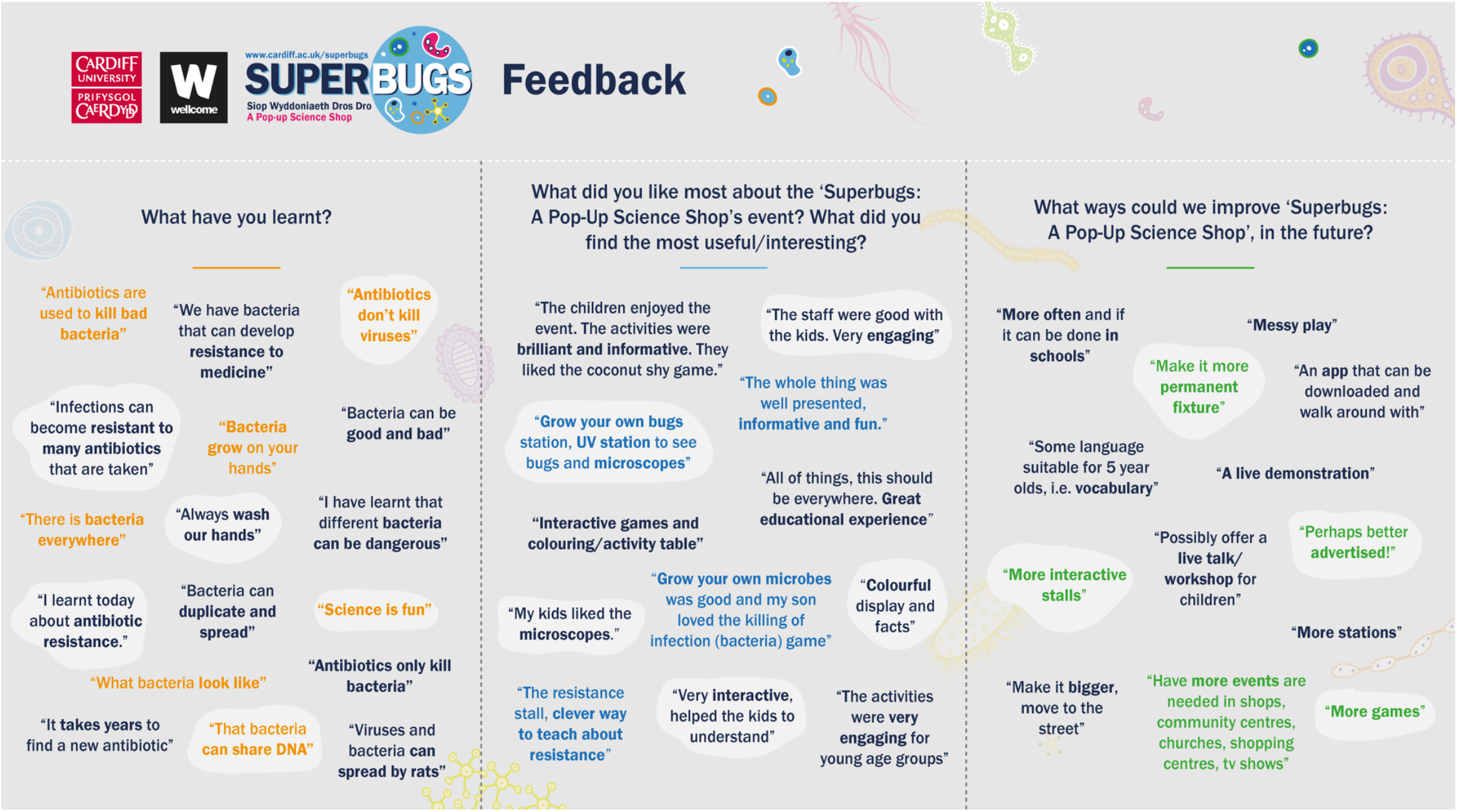
Word clouds summarising feedback received by visitors. **(A)** Summary of the feedback received from 116 visitors on the ‘bacterial thought’ tree. (B) Summary of 202 responses: “What did you like most about the ‘Superbugs: A Pop-Up Science Shop!’ event? What did you find the most useful/interesting?”. **(C)** Summary of 53 responses to: “What ways could we improve ‘Superbugs: A Pop-Up Science Shop!’, in the future?”

**Figure 6:**
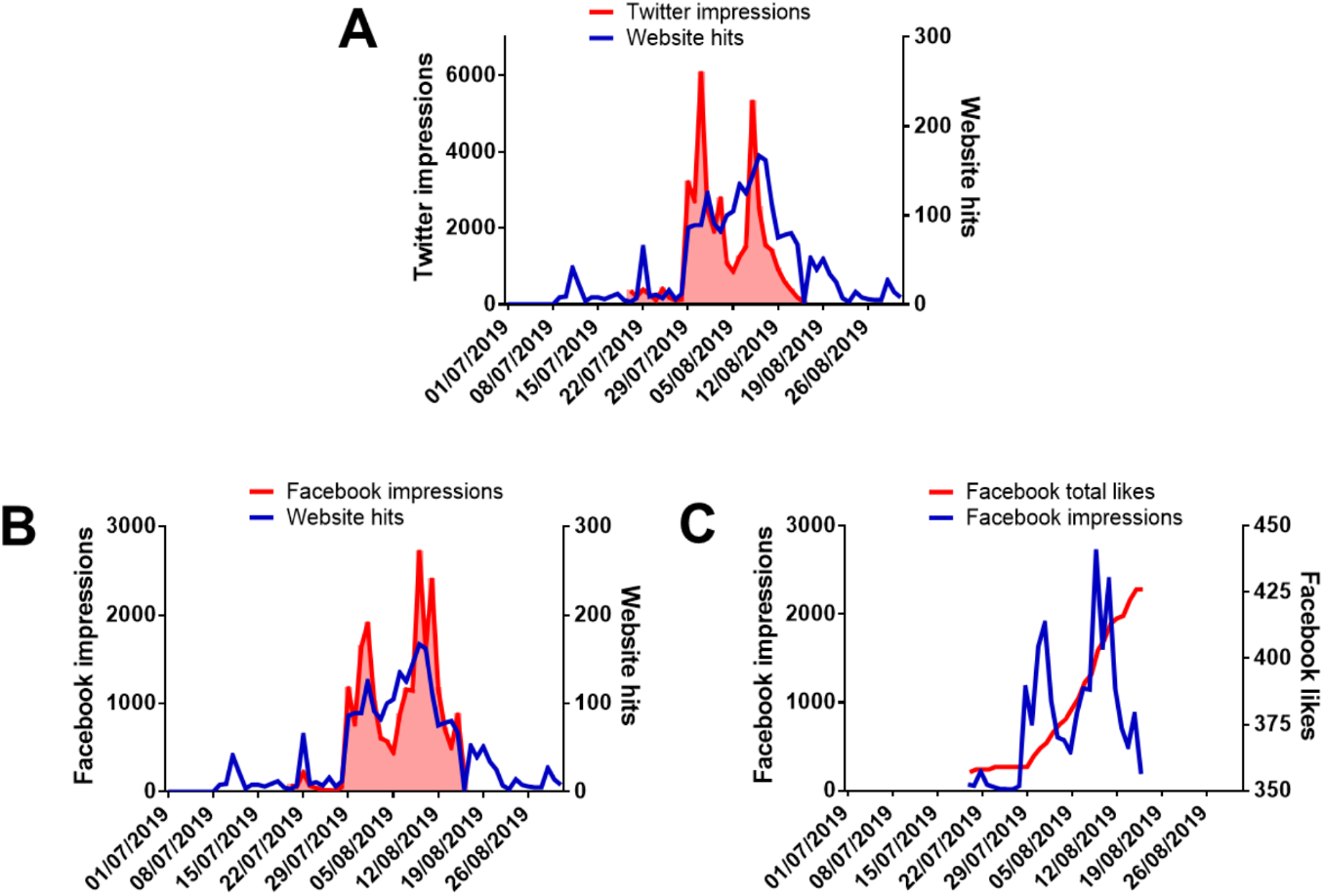
Social media activity and web traffic before, during and after the ‘Superbugs’ event. **(A)** Time course of direct hits on the ‘Superbugs’ website (www.cardiff.ac.uk/superbugs/) and impressions on the @CUSystemsImmu Twitter account; **(B)** Time course of direct hits on the ‘Superbugs’ website and impressions on the SystemsImmunity Facebook account; **(C)** Time course of impressions on the SystemsImmunity Facebook account and development of Facebook followers.

### 6.3 Antibiotic Resistance Champions

At Station 1 each young visitor was given a sticker card, corresponding to 6 different activities throughout the shop (Stations 4, 5, 6, 8, 9 and 11; Figure 4). Once all 6 stickers were collected they were awarded the title of Antibiotic Resistance Champion, their choice of ‘Superbugs’-branded prize (pen, bookmark, badge or balloon) and a certificate with handy tips of how they can help in the fight against AMR (Figure 7). We created 1,626 Antibiotic Resistance Champions, from across 200 different schools in Wales, and many further afield (Figure 7). This approach adopted the theories underlying the ‘behavioural pledge’ approaches that were previously successful in handing a level of responsibility to stakeholders to facilitate positive actions and to reinforce their aims and objectives (Kesten *et al*., 2017; Little *et al*., 2015; Eley *et al*., 2018).

**Figure 7:**
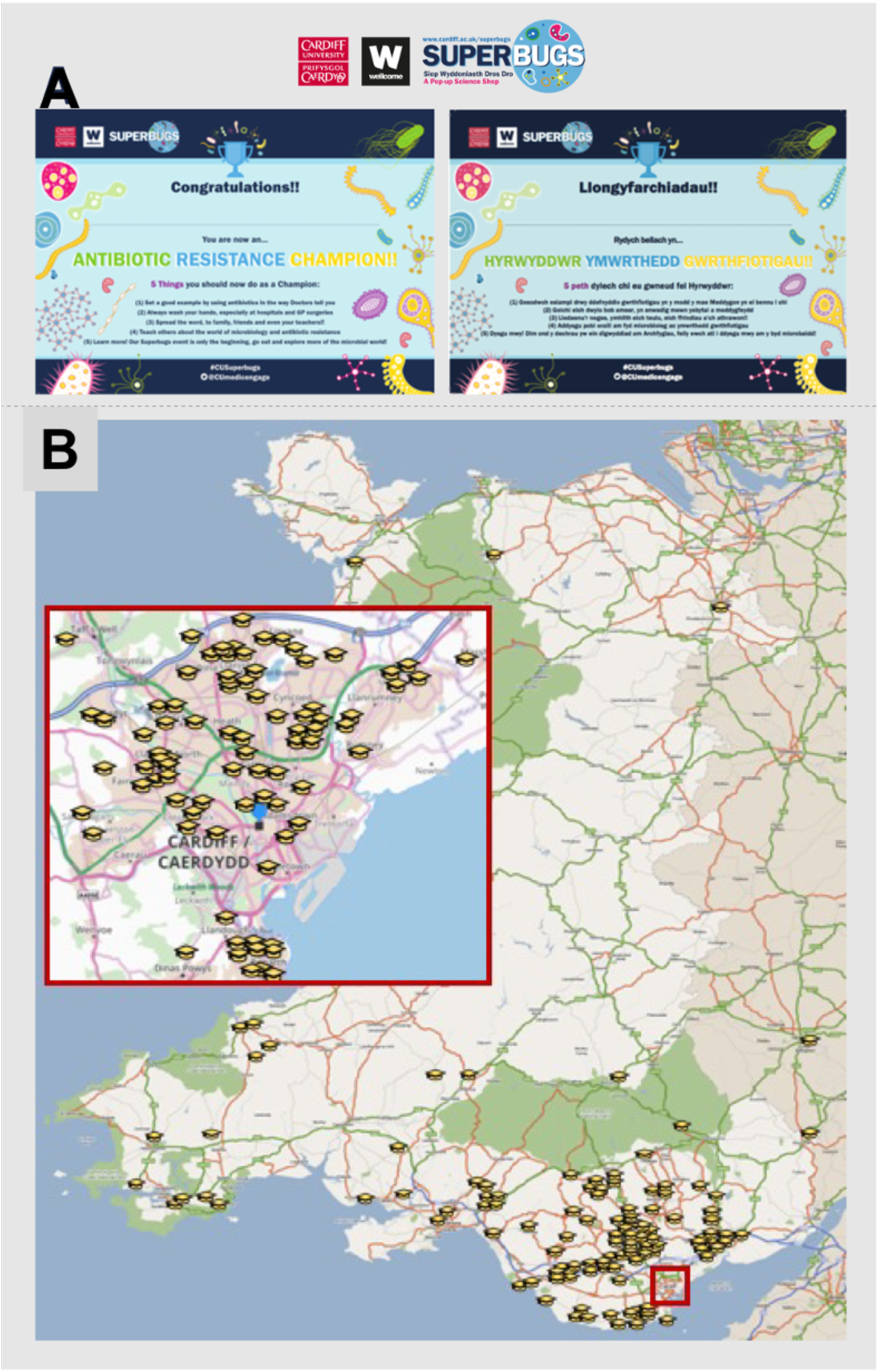
Antibiotic Resistance Champions: (**A)** Bilingual Antibiotic Resistance Champion certificate; **(B)** Distribution of schools at which the new Antibiotic Resistance Champions attended at the time of the event. Other Champions were created in England and internationally (France, Belgium, Spain, Jersey Channel Islands and Kuwait).

### 6.4 External Outputs

The event received a notable amount of local media attention. We were involved in a fun feature on the ‘Josh & Kally show’ for Capital FM South Wales. The radio presenters were invited to swab themselves live on air, and we returned the next day to deliver the results as to who grew up the most microbes. This was a unique way to engage a young demographic in the topic of bacteria and to inform wider groups about the ongoing ‘Superbugs’ event. The local television station Cardiff TV spent over an hour in the shop filming visitors engaging with our activities, and interviewing members of the ‘Superbugs’ team. Two videos were produced and shown repeatedly across the local area for the remainder of the event and can be found at the events homepage (https://www.cardiff.ac.uk/systems-immunity/engagement/understanding-science/superbugs-a-pop-up-science-event).

We were also able to raise interest and attention from high profile stakeholders including Kirsty Williams AM (Welsh Minister for Education & Skills) and a visit from the local Member of Parliament for Cardiff Central, Jo Stevens (the current Shadow Secretary of State for Digital, Culture, Media and Sport), in addition to tweets/retweets by a number of Welsh journalists. Jo Stevens took part in a number of ‘Superbugs’ activities, including swabbing her mobile phone to grow any contaminating microbes, and wrote a blog about her experience.

More recently, ‘Superbugs: A Pop-up Science Shop!’ was accepted as an entry to the National Co-ordinating Centre for Public Engagement (NCCPE) ‘Engage 2020’ conference that took place online from 30th November – 4th of December 2020. Our video submission can be found on YouTube in NCCPE’s ‘Examples of Practice’ playlist (https://www.youtube.com/watch?v=NuPnWqz8FdU&t=1s).

## 7. Impact of Project

### 7.1 Impact on Public Engagement Delivery Strategy

A primary aim of ‘Superbugs: A Pop-Up Science Shop!’ was to evaluate this form of public engagement, a novel scheme for both the School of Medicine, and Cardiff University as a whole. We hypothesised that this mode of delivery would be successful in reaching a wide demographic, beyond the limited cohorts who would typically seek out opportunities to engage in scientific research. Questionnaire data collected showed that indeed 67.3% of entries were impromptu visits with no prior knowledge of the event before spotting it on the concourse of StD (Figure 8). This was significantly higher than those visitors indicating any awareness due to aspects of our promotional campaign, including advertisement in the Primary Times magazine, website and social media. Whilst confirming our hypothesis, it was also a triumph for the imaginative and imposing exterior designed by our team and informed by our focus group. This also suggests that in future public space located engagement events, significant resources may be better focused on providing the best and most attractive possible experience for visitors, as opposed to an overly enthusiastic promotional campaign.

**Figure 8:**
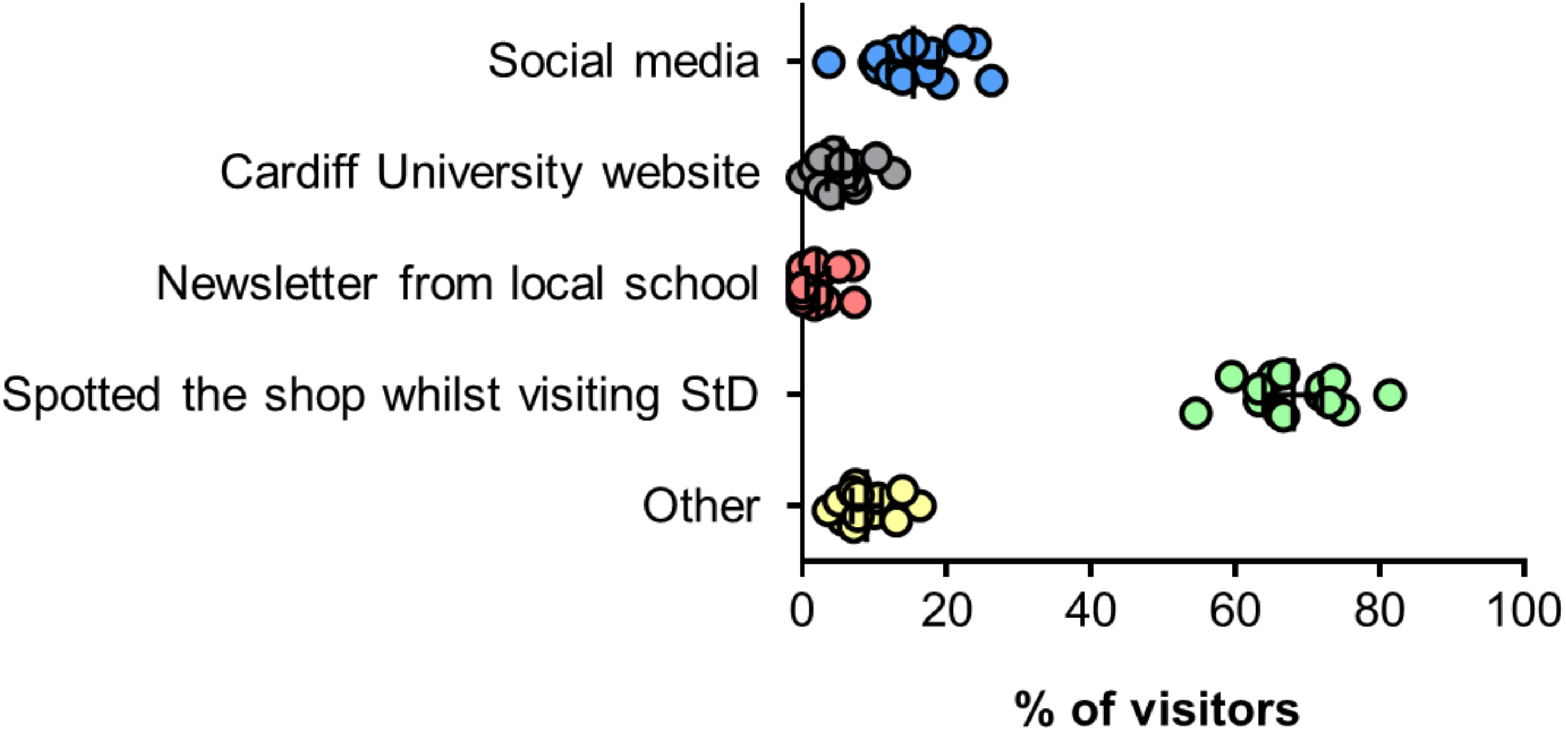
‘Where did you hear about Superbugs: A Pop-up Science Shop!’ [Questionnaire Q9]. **(A)** Social media (Facebook, Twitter), including Primary Times; **(B)** Cardiff University website; **(C)** Newsletter from local school; **(D)** Spotted the shop whilst visiting StD; **(E)** Other. Data collected from questionnaires over 14 consecutive days (n=656 in total). Individual data points show daily answers; error bars show mean values and 95% confidence intervals.

**Figure 9:**
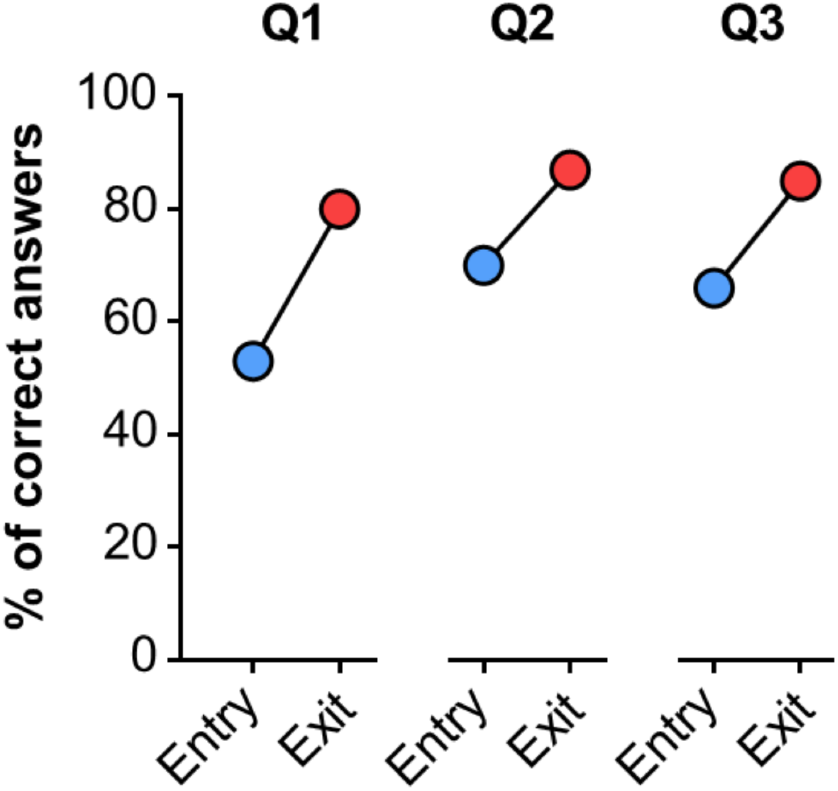
*Comparison of correct answers to benchmarking questions by Stakeholders on Entry and Exit to ‘Superbugs’* Benchmarking questions are listed below with correct answers underlined. **[Q1]** “Antibiotics are used to kill…..” (A) Viruses, (B) <u>Bacteria</u>, (C) Fungi, (D) All of the above. **[Q2]** “When taking antibiotics you should…..” (A) stop when you feel better, (B) <u>Take all the antibiotics as instructed by your doctor</u>, (C) Save some for next time you feel unwell, (D) Share them with your friends. **[Q3]** “Antibiotic resistance is a problem for…..” (A) only people taking antibiotics, (B) <u>Everyone</u>, (C) The elderly & sick, (D) Those who travel to exotic countries.

The importance of our focus group in shaping the nature of the content and the overall design theme for ‘Superbugs’, and the impact this then had on the success of the event, illustrates the potential in utilising focus groups, public involvement and co-production in shaping such projects. We hope this leads to developing a culture of increased efforts for co-production and public involvement across the School of Medicines engagement activities, to the benefit of the scientists and public alike.

Further confidence in our strategy was corroborated in the data collected through our questionnaires (Table 2). 95% and 91.9% of visitors, respectively, agreed/strongly agreed that the event was fun, engaging and informative, and that we had pitched the intellectual level appropriately. We are very proud to say that over 94% of visitors indicated that they not only rated ‘Superbugs’ as ‘Very Good’ or ‘Excellent’ but, perhaps even more significantly, they would also recommend ‘Superbugs: A Pop-Up Science Shop!’ to others.

**Table 2:** Percentage of stakeholders that selected ‘Agree’ / ‘Strongly Agree’ with the below statements. Data collected from questionnaires (n=656).

|  | Strongly Disagree | Disagree | Neutral | Agree | Strongly Agree |
| --- | --- | --- | --- | --- | --- |
| [Q1] “Superbugs was a fun, engaging, informative event” | 3.7 | 0.2 | 1.1 | 27.9 | 66.7 |
| [Q2] “We have a better understanding of the world of microbes because of Superbugs” | 3.8 | 0.3 | 3.1 | 38.4 | 54.3 |
| [Q3] “We have a better understanding of antibiotics and how they work” | 3.9 | 0.3 | 3.3 | 37.5 | 55.0 |
| [Q4] “We have a better understanding of antibiotic resistance” | 4.0 | 0.6 | 3.7 | 35.4 | 56.3 |
| [Q5] “We are more likely to take a full course of antibiotics” | 3.9 | 0.8 | 4.5 | 33.4 | 57.4 |
| [Q6] “We are likely to tell others about what we have learnt” | 4.1 | 0.6 | 4.1 | 33.5 | 57.8 |
| [Q7] “The intellectual level of content was pitched appropriately for a family audience” | 3.9 | 0.6 | 3.6 | 33.6 | 58.2 |
| [Q8] “We would recommend ‘Superbugs: A Pop-Up Science Shop!’” | 4.0 | 0.2 | 1.4 | 19 | 75.4 |

Following a visit, Professor Kim Graham, Pro-Vice Chancellor for Research, Innovation and Enterprise at Cardiff University and Professor Gary Baxter, Pro-Vice Chancellor for the College of Biomedical and Life Sciences at Cardiff University took to social media with positive tweets of their experience and encouraged others to visit the shop and event. This engagement from senior leaders at the University highlighted the high-level support for the Superbugs project and public engagement with research. Future conversations were invited from senior leaders, to explore this model of engagement, its potential for societal change and how such activities can be sustained in the longer term.

### 7.2 Impact on AMR Awareness for Stakeholders

We may assuredly accept the hypothesis that our pop-up shop was an effective way of imparting positive impact on the AMR awareness of the public audience with whom we engaged. Table 2 illustrates that ‘Superbugs’ improved the understanding of the world of microbes, antibiotics (and how they work) and antibiotic resistance in 92.7%, 92.5% and 91.7% of cases, respectively. Perhaps more significantly, our evidence supports a progressive influence on future behaviour and attitudes towards antibiotic stewardship, with 90.8% of participants now more likely to complete a full course of antibiotics. Improved antibiotic stewardship has significant and far-reaching implications for our ability to control AMR pathogens (File *et al*., 2014)

Secondary to this, we may infer far-reaching legacy impact of our event. 1,626 young people left not only bestowed with the title of ‘Antibiotic Resistance Champion’ but also with their certificate detailing tips on how they can further their knowledge and spread the word long after the doors of ‘Superbugs’ had closed. As a testimony to the success of this approach, 91.3% of visitors completing the questionnaire stated they would pass on what they had learnt at the event (Table 2).

### 7.3 Impact for the Institute and Staff

‘Superbugs’ provided a valuable opportunity for professional development for all involved. For the lead scientist, it was their first experience of independently securing grant funding, and the organisational and administrative responsibilities that comes there-in.

Logistically, ‘Superbugs’ was a mammoth undertaking requiring staff to give up their own time to take part. In total, 33 volunteers helped deliver aspects of the event, involving 5-6 individuals being present at the shop and facilitating the activities at any one time. Station 4 (‘Grow Your Own Microbes’) in particular required extra-curricular assistance in the incubation, sorting and photography of all swab plates, and the subsequent uploading of the anonymised photos onto a bespoke Facebook album. Volunteers were at various stages of career and backgrounds across the academic spectrum; from professional staff, research associates and senior academics, to students ranging from undergraduates to PhD, and for many ‘Superbugs’ represented their first experience of public engagement. On their first day of volunteering, all were fully briefed on the content of the shop, and the concept and messages behind each activity. Each day we attempted to provide a mixture of inexperienced ‘engagers’ with more senior staff, in order to breed a supportive environment where communication skills could be developed and enhanced.

As described, we saw an incredible response to our Station 5 ‘Grow your own microbe’ activity, where visitors were able to swab themselves and then using a unique code, follow up on what grew at a later date online. The enthusiasm with which visitors engaged with this suggests at a potential rich resource that could be exploited for research purposes and could provide a unique insight into the social and environmental spread of target micro-organisms, and the anthropogenic factors driving this dissemination.

‘Superbugs’ instigated a significant influx of activity and attention onto the social media and web pages of the School of Medicine public engagement team. Across the academic year, the School of Medicine delivers a wide portfolio of engagement activities for students of primary and secondary schools, and the wider public, including a free-access Public Lecture Series, the Wales-wide Life Sciences Challenge quiz and the ‘Science in Health Live!’ event and laboratory work experience scheme which both provide unique opportunities for year 12 school pupils to visit the working environment of the University Hospital of Wales in Cardiff (https://www.cardiff.ac.uk/medicine/about-us/engagement) witnessing cutting edge technologies and research first hand.

### 7.4 Access for protected characteristics

A cornerstone to the delivery strategy of ‘Superbugs’ was the hope to breakout out from traditional cohorts of those scientifically aware, to wider and more diverse demographics, including those of age, culture and religion. Enrolling Diverse Cymru as a strategic partner provided us with an independent and judicious evaluation on how ‘Superbugs’ met the needs of, and appealed to, people with protected characteristics, and ways in which we could improve this even further for future activities. This was considered during the design of the shop layout, ensuring information and activities were accessible for a range of heights and abilities, and there was adequate space to accommodate mobility aids. Highlighted was the need to more apparently cater for those with sensory, learning and cognitive impairments, primarily in some of the language used at certain areas of the shop, and the prominence of instructions for our activities. In addition to this, we are committed to applying decolonisation of all information provided in future ‘Superbugs’ events, to reflect a more accurate, diverse and global view of the topic area.

## 8. Discussion

### 8.1 Summary of Principal Findings

To our knowledge this is the first pop-up science shop designed to increase the awareness and knowledge of the general public of the microbial world, infection biology and the increasing threat of antibiotic resistance to global public health. Attracting 6,566 visitors in a two-week period during the summer school holidays indicates a strong thirst for such engagement in research by members of the public.

Based on the qualitative and quantitative data collected, the anecdotal feedback received, and personal communication from senior academic colleagues across the School of Medicine, we are confident in concluding that ‘Superbugs: A Pop-up Science Shop!’ made positive strides in raising awareness of the AMR crisis and educating the public of the part they need to play in the fight against this global issue. The data presented herein indicates that we have met the aims we set and achieved the outputs we had strived for at the beginning of the ‘Superbugs’ project. We created an environment of two-way dialogue with wide and varied public demographics, imparting a positive change in the awareness of microbiology and AMR. Concurrently, ‘Superbugs’ has illustrated the efficacy of public space-based engagement in doing so.

### 8.2 Limitations and Future Improvements

‘Superbugs’ was delivered in one city centre location, Cardiff (Wales’s capital). It is the ambition of the project team that future redeliveries of ‘Superbugs’ events would act to widen the accessibility to include other cities and towns across Wales, particularly those outside of southern hubs. This would enable us to compare the impact of ‘Superbugs’ across different geographical and socio-economic areas, of more diverse stakeholder demographics, and provide deeper data sets with which to evaluate the efficacy of our engagement model.

As has already been indicated, this project was built on upon the time and hard work of the core project team, and the willingness of an army of volunteers to give up their own time to participate. In the lead-up to the event itself, we begun a recruitment drive through the communication networks of the School of Medicine, and more widely the College of Biomedical Life Sciences. Regrettably, uptake outside of the research groups of the involved academics was limited, and as such at certain times numbers during the event were limited to the point of threatening to compromise the experience of our visitors. Notwithstanding, ‘Superbugs’ presented a valuable, but for many a missed, opportunity to develop communication and outreach skills, and gain insight into how public engagement activity with research may support the development of pathway to impact statements. It should be noted that one potential contributor of this was the timing of the event during the summer holidays, perhaps suggesting the importance of budgeting of staff time in the organisation of large-scale engagement and outreach events.

This also highlights an intrinsic issue within the academic culture and attitudes towards public engagement activities. Historically, public engagement may be seen simply as a way in which to educate the ‘scientifically illiterate’. As such, the discipline of public engagement has been underserved in time, attention, funding, and willingness to participate (as evidenced here) by academic institutions and the staff therein. Perhaps nothing more than a pleasantry aside to the primary roles of researchers. This is further exacerbated by an environment that prioritises quantifiable publications and grant funding in determining career prospects and progression. This apathy is somewhat counterintuitive given all science, at it’s very foundation, is for the betterment of the human condition and those we seem reluctant to communicate to.

The introduction of ‘impact’ (encompassing public engagement) as an element of assessment for REF 2014 is perhaps a clear signal of a slowly changing tide in this regard (Copley, 2018). Increasingly, public engagement may be seen as a tool to raise an institutes profile, influence policy makers (both directly and indirectly) and further quantify research impact. However, a paradigm shift in the way in which public engagement with research is regarded at all levels of academia is still needed. More standardised frameworks for the design, implementation and evaluation of public engagement activities are required to advance the integrity and rigour of this capacious discipline, and there is a growing literature to achieve this (Mahony and Stephansen, 2017). Furthermore, as the demand for public engagement with research activities continues to increase from funders, concern around recognition, value and support amongst employers requires further exploration.

## 9. Conclusion

Currently, we as a society are facing the challenges of the COVID-19 pandemic. It would be remiss however, to entertain any oversight of the more silent pandemic of AMR that has been with us for decades and continues to carry severe impact economically and on public health. Increasingly, a consensus view is emerging that the COVID-19 pandemic actually threatens a further exacerbation of the antibiotic resistance crisis worldwide (Miranda *et al*., 2020; Murray, 2020). An alternative view holds that the reduction of international travel may prohibit the global dissemination of AMR pathogens, and scientific communication regarding this viral pandemic may infer improved awareness around the appropriate use of antibiotics (Murray, 2020). It is clear then that there will be a post-COVID-19 impact on AMR, even if the true nature and parameters of these are not yet known (Monnet and Harbarth, 2020). Either way, it is evidently pertinent to continue effective and impactful engagement of the public on the topic of infectious disease.

‘Superbugs’, which is now an inter-institutional collaboration between academic staff of the University of Bristol and Cardiff University, presents unique opportunities for the project’s future in terms of size, scope, and further funding. As with all things, ‘Superbugs’ has had to adapt to a new COVID-19 world where large scale in person engagement events are simply not possible at present, and not for the foreseeable future. In August 2020 we successfully secured grant funding through the ISSF3 Public Engagement Co-production award to co-produce a permanent online/digital website presence for ‘Superbugs’ and are currently working with stakeholders from across the education sector, whilst designing and delivering our co-production and evaluation strategies. Immersive and interactive events in public spaces will continue to be at the heart of what we deliver with ‘Superbugs’, and we were pleased that just before the arrival of COVID-19 in Wales, a first successful redelivery of select ‘Superbugs’ activities were run as part of Cardiff Science Festival in February 2020.

Increasing public awareness around AMR and antibiotic use now forms a cornerstone of the UK government’s 20-year strategy in managing and controlling the issue (www.gov.uk/government/publications/uk-20-year-vision-for-antimicrobial-resistance).

Redfern *et al*., (2020) comprehensively outlined a number of activities, representing a diverse range of approaches that have been implemented to this end, and in doing so highlights a most pertinent point. Simply, in undertaking a systematic review of AMR-related engagement activities, an intrinsic limitation was the relative paucity of publications based on such activities. Indeed, there is a similar such problem for microbial literacy within the public and education sectors also (Timmis *et al*., 2019) and this is further reflected in the Wellcome Trust’s ‘Reframing Resistance Report’ (https://wellcome.org/sites/default/files/reframing-resistance-framing-toolkit.pdf). We hope that accounting ‘Superbugs: A Pop-Up Science Shop!’ herein we will contribute to a growing body of work laying the foundations to address such a problem, and in doing so leave a lasting impact on addressing public awareness of the AMR threat, and the best practices to achieve this aspiration.

## 10. Notes on Contributors

**Dr Jonathan M. Tyrrell** is a Lecturer in Medical Microbiology and Antibiotic Resistance at the School of Cellular & Molecular Medicine, University of Bristol. As a passionate scientific communicator Jonathan became involved in a number of engagement activities, leading to him creating ‘Superbugs: A Pop-Up Science Shop’. Jonathan was lead scientist on the ‘Superbugs’ and was responsible for the conception, and involved in all following facets, of the project.

**Ms Christie Conlon** is a professional graphic designer for Cardiff University’s College of Biomedical & Life Sciences and the BBC. Christie was the graphic artist of the project and was responsible for the design of the shop interior, exterior, logos, certificates and give-aways produced by the project.

**Dr Ali F Aboklaish** is a research associate working on antibiotic resistance in the School of Medicine, Cardiff University. Ali assisted in the administrative, planning and delivery of the project and was also a member of the ‘Science content’ team that designing the information and activities delivered.

**Mrs Sarah Hatch** is Cardiff University’s School of Medicine’s Engagement Manager. Sarah led on the organisation of the project’s focus group ensuring the project’s target audience informed the development, delivery and subsequent output of the proposed engagement activity. Sarah also took a significant role in developing the evaluation strategies of the project.

**Mr Carl Smith** is the Public Engagement Manager at Cardiff University. Carl was an advisor from the earliest stages of the project and volunteered for the delivery of the event.

**Mr Jordan Mathias** is a research assistant working in the field of antimicrobial resistance in the School of Medicine, Cardiff University. Jordan was part of the ‘Science content’ team, designing the information boards and activities within the shop. Jordan has now joined the core team, helping to deliver the next stages of our ‘Superbugs’ project.

**Ms Katy Thomson** is a PhD student working in the field of antimicrobial resistance in the School of Medicine, Cardiff University. Katy was part of the ‘Science content’ team, designing the information boards and activities within the shop.

**Professor Matthias Eberl** is Professor of Translational Immunology and Academic Lead for Public Involvement and Engagement at the School of Medicine, Cardiff University. Matthias was instrumental in the strategic design, delivery and evaluation of ‘Superbugs’.

## 11. Acknowledgements

We would like to thank all volunteers for their endless efforts in supporting the project from conception to delivery, without whom the project would not have been possible (in alphabetical order): Raya Ahmed, Abdullah Almusallam, Sarah Baker, Rebecca Bayliss, Ian Boostrom, Simone Cuff, Karen Edwards, Refath Farzana, Ana Ferreira, Hyun-Sun Jin, Giulia Lai, Mei Ling, Mark Ponsford, Edward Portal, Savitha Radhakrishan, Jacob Rees, Georgina Russell, Mike Roberts, Kirsty Sands, Louise Stack, Mark Toleman, Lucas Tselepis, Timothy Walsh, and Janis Weeks, Qiu Yang. A special thanks must be extended to Marie-Claire Bell and Emma Widlake. Further thanks must go to Thomas Greaves of Morgans Consult, whose support in delivering the installation of the exterior and interior design and fitting of ‘Superbugs’ was unwavering and beyond the call of duty. A thank-you to the British Society for Immunology who provided important support to our grant application and to the event. Finally, a thank you to GiantMicrobes^®^ for their generous provision of cuddly microbes to be used as display items and prizes to be won.

